# Identifying foraging spatial cues in beehive sound activity using machine learning methods

**DOI:** 10.1101/2024.07.08.602596

**Authors:** David Bocanegra, Jorge Galvez, Nayan Di, Fanglin Liu, Fernando Wario

## Abstract

The beehive sound, a continuous signal produced by bees within the hive, has been found to correlate with different behavioral states of the colony, like being queenless and swarming. We investigated the possibility of identifying foraging spatial cues in this signal. We recorded a colony’s sound while foraging from food sources located at three different distances from the hive, one at a time. The recordings were split into frames to obtain six statistics of their Mel Frequency Cepstral Coefficients. Then, we evaluated different autoencoding networks to obtain a latent space that allowed frames from different foraging distances to be easily differentiable. The high accuracy, silhouette score, and F1 score shown in the obtained latent spaces strongly support our approach for identifying foraging spatial cues in beehive sound activity.

## 1 INTRODUCTION

The honey bee waggle dance is one of the prime examples of abstract communication in the animal world. Worker bees use it to communicate to their nest mates the location of resources such as nectar, pollen, and water. Foragers who return from a valuable food source move in a stereotypical pattern on the honeycomb surface, and some of the bees that closely follow their movements are recruited to the source location (von Frisch, 1965).

During the waggle dance, the dancer vigorously shakes her abdomen while moving forward in what is known as the waggle phase of the dance. At the end of each waggle phase, the dancer circles back to the starting point, alternating right and left, tracing a trajectory that resembles the number eight. Each dance may consist of dozens of waggle-return phases, with the number of cycles and the overall duration of the dance correlating with the source profitability (Seeley et al., 1991, 2000).

To the human observer, the waggle phase of the dance can be interpreted as a vector encoding the polar coordinates to the food source (see *Figure 1*). As demonstrated by Karl von Frisch, the dancer’s body orientation relative to the hive vertical during the waggle phase reflects the direction of the food source relative to the sun’s azimuth in the field. Complementary, the duration of the waggle phase correlates with the distance to the food source (Gould et al., 1970; Riley et al., 2005; von Frisch, 1965).

**Figure 1.**
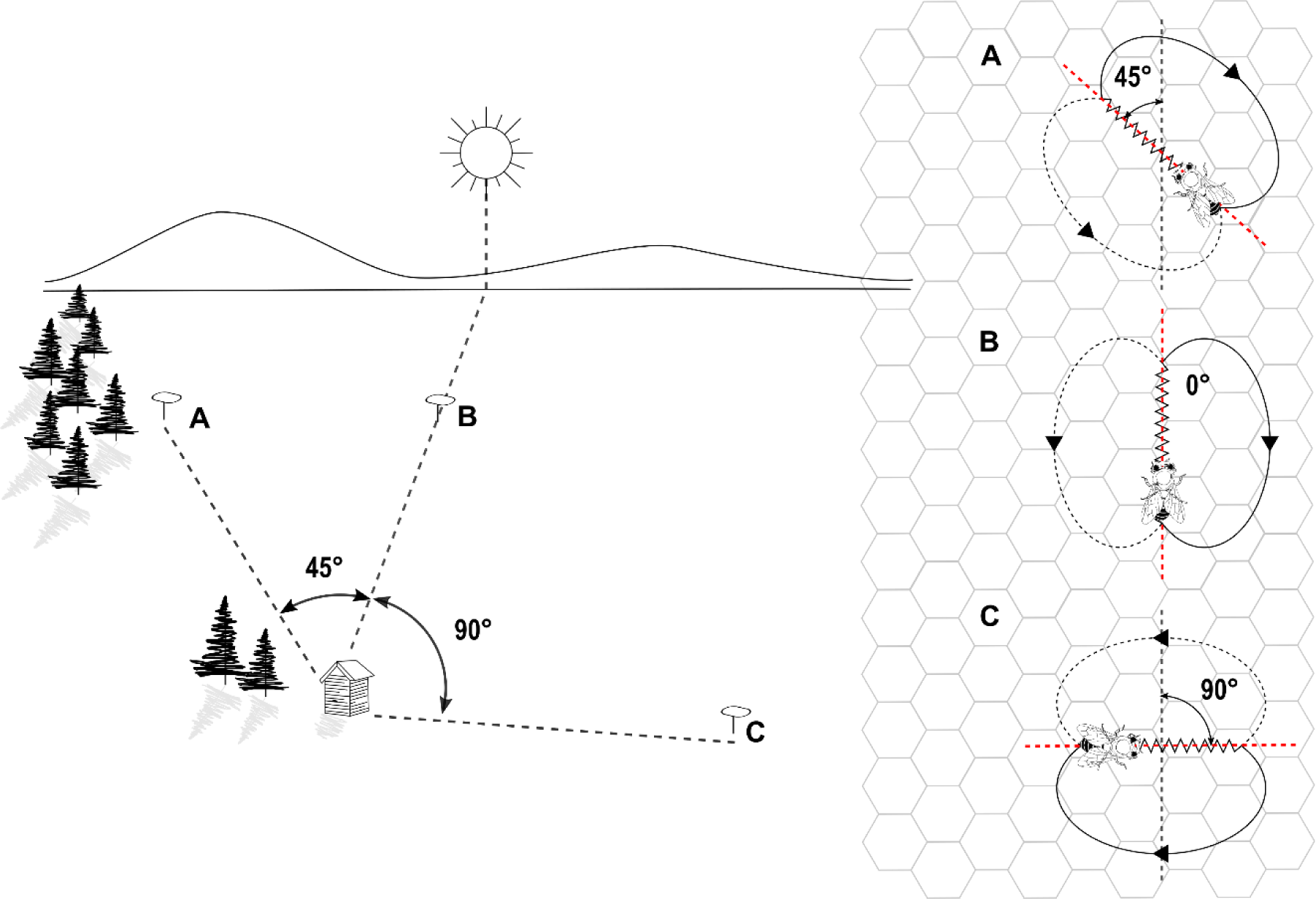
Correlation between waggle dance parameters and locations on the field. On the left side, three food sources in the field located at A) 45° counterclockwise, B) 0°, and C) 90° clockwise relative to the solar azimuth. On the right side, the representation of food sources A, B, and C through waggle dance paths on the surface of a vertical comb. Figure available under CC BY 4.0 from (Wario et al., 2017).

The repeated cycles of waggle and return phases endow the waggle dance with valuable redundancy, although always limited to the same communication channel. Several studies indicate that along with the visual cues, pheromones, substrate vibrations, and airborne sounds (Kirchner, 1993; Michelsen et al., 1989; Tautz, 1996; Thom et al., 2007) also play a role in the waggle dance, specifically in attracting and recruiting followers. During the waggle phase of the dance, bees emit an airborne pulsed sound at the frequency of 250-300 Hz (Michelsen et al., 1986; Wenner, 1964), a signal that could function as an additional communication channel for redundancy in the waggle dance.

Recording the visual components of the waggle dance can be fairly unintrusive thanks to the observation hive, which provides visual access to virtually all activity happening on the comb surface, allowing the development of automatic techniques for the study of dance communication (Bozek et al., 2021; Schürch et al., 2016; Wario et al., 2015, 2017). In contrast, recording a dancer’s sound and pheromone emissions, due to their nature and the high population density on the comb surface, is a complex task that usually requires operating an open hive (Thom et al., 2007; Wenner, 1964), which, in turn, may disturb the colony and introduce high noise levels to the data.

Given this challenging scenario, the study of pheromones and sound signals is usually conducted at the colony level. The “beehive sound”, or “colony noise” is the collective buzzing and vibrating sound produced by the activity of honey bees within the hive. Characterized as a series of low-frequency vibrations and higher-frequency buzzing noises, the beehive sound has been proven to strongly correlate with the colony size, activity level, and environmental conditions (Kirchner, 1993; Sharif et al., 2020; Terenzi et al., 2020; Yu et al., 2022).

Audio-based classification models have already been applied to beehive sound to detect key statuses of the colony, such as being queenless (Cejrowski et al., 2018; Howard et al., 2023), swarming (Ferrari et al., 2008), and the presence of diseases or pests (Robles-Guerrero et al., 2017). A variety of classification approaches have been proposed for this task, including Principal Component Analysis (PCA), Support Vector Machines (SVM), Linear Discriminant Analysis (LDA), Convolutional Neural Networks (CNN) (Nolasco & Benetos, 2018; Qandour et al., 2014; Rybin et al., 2017), and Random Forest (Robles-Guerrero et al., 2017).

Classification models do not operate directly on the audio amplitude time series but rather on acoustic feature sets. Three feature sets have proved to work for the classification of beehive audios, namely low-level signal features, including Zero-Crossing Rate, Bandwidth, Spectral Centroid, and Signal Energy (Qandour et al., 2014), Mel Frequency Cepstral Coefficients (MFCC) (Nolasco & Benetos, 2018; Robles-Guerrero et al., 2017), and soundscape indices (Mammides et al., 2017; Sharif et al., 2020).

In this work, we propose the use of a classification model to explore the presence of foraging spatial cues in the beehive sound. The success of the classification model relies on a proper combination of features and classification methodology. Inspired by the work of Scarpiniti et al. (Scarpiniti et al., 2021a), where an audio classifier for environmental sound is successfully applied in the challenging scenario of construction sites, we also decided to use a statistically optimized set of features obtained from Mel-frequency cepstral coefficients (MFCCs) (Robles-Guerrero et al., 2017). As a classifier, we use an artificial neural network which is first pre-trained as an autoencoder. The pre-training process helps to initialize the classifier’s parameters in more promising areas of the parameters space, which after fine-tuning results in better generalization (Erhan et al., 2010).

The positive results for identifying the foraging spatial cues in beehive sound validate the use of autoencoding networks in combination with MFCC’s statistical features as a methodology for classifying sound signals. To the best of our knowledge, this is the first time that foraging spatial cues have been successfully identified in beehive sound activity, calling for further studies in this direction that could improve our understanding of sound as a means of communication in honey bee colonies.

## 2 MATERIALS AND METHODS

### 2.1 Audio data

The experiments were conducted at the Sericulture and Apiculture Research Institute of Yunnan Academy of Agricultural Sciences in Mengzi City, Yunnan Province, China (23.52°N, 103.39°E) between November 2020 and January 2021. The colony of the species Apis Cerana resided in a typical wood beehive with a 10-month-old queen, was healthy and without any signs of attack by pests, emerging diseases, or viruses. At the beginning of the first week, foragers from the colony were trained to collect sugar water from an artificial feeder 100 m from the hive entrance. After **7** days of foragers visiting the feeder, the beehive sound is recorded at the time of the highest affluence with a microphone located inside the beehive at 15 cm from the floor. The process was repeated in the following weeks with the feeder located in the same direction at 300 m, and 500 m from the hive.

The microphone used for the audio recording was a PCK200 from Takstar, with a frequency response of 30 Hz to 20 kHz and a sensitivity of -36dB±3dBV. The microphone was connected to a digital sound card UM2 from Behringer and processed using the software Audacity at a sampling frequency of 44,100 Hz and a bit rate of 1,411 kbps in mono. Finally, the audio files were saved in the WAV format, with each file covering over 40 minutes of beehive sound activity (see *Table 1*).

**Table 1.**
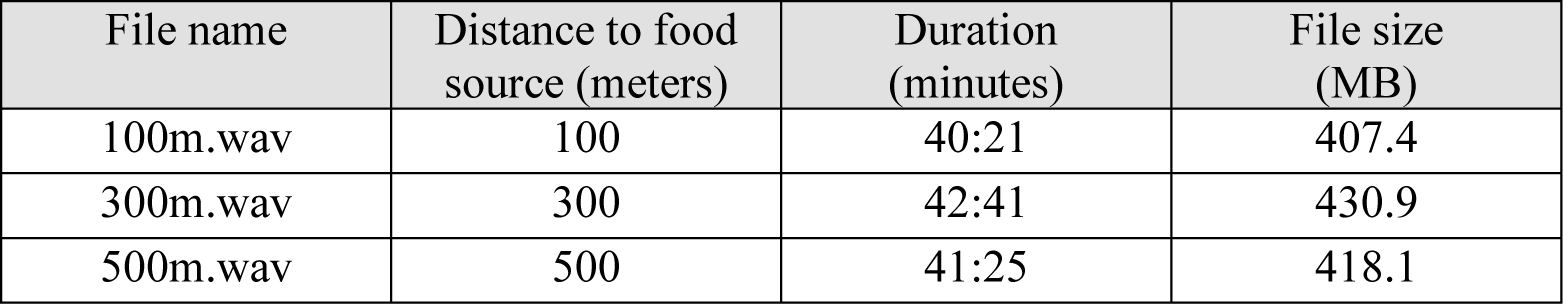
Information on the audio recordings for the three experiments considered in the study.

| File name | Distance to food source (meters) | Duration (minutes) | File size (MB) |
| --- | --- | --- | --- |
| 100m.wav | 100 | 40:21 | 407.4 |
| 300m.wav | 300 | 42:41 | 430.9 |
| 500m.wav | 500 | 41:25 | 418.1 |

### 2.2 Autoencoders

Autoencoders (Rumelhart & McClelland, 1987) are a type of neural networks trained in an unsupervised manner to reconstruct their input data. An autoencoder consists of two subsystems, an encoder and a decoder. The encoder works as a function *f* that maps an input *X* ∈ ℝ^*N*^ to an encoded representation (called latent representation) *H* ∈ ℝ^*M*^ (Equation

(1)), while the decoder operates as a function *g* that generates an approximation *X*′ ∈ ℝ^*N*^ of the input *X* from the representation *H* generated by the encoder (Equation
(2)). To find a proper set of functions [*f*, *g*], the autoencoder is trained using a loss function, like the Mean Squared Error (MSE), that measures the difference between the reconstructed data *X*′ and the input data *X*. Figure 2 shows a graphical representation of an autoencoder.

**Figure 2.**
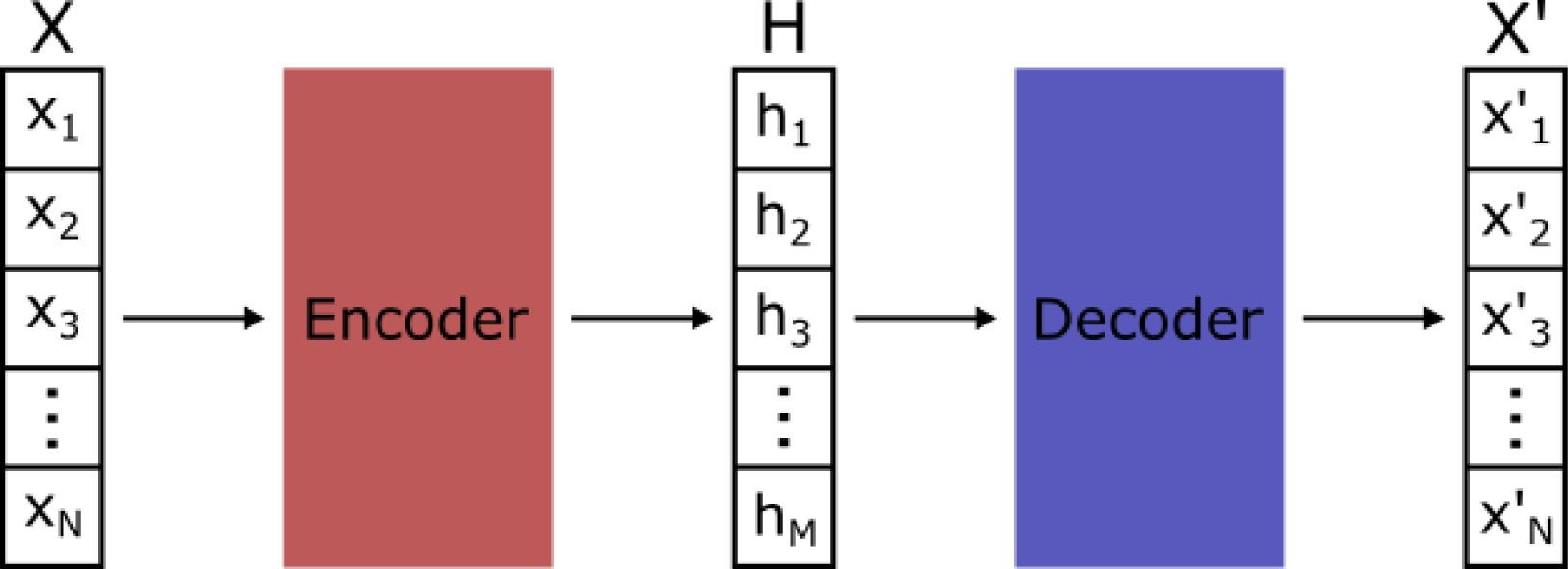
Graphical representation of an autoencoder. First, the input vector X of dimension N is mapped by the encoder to a vector H of dimension M, then the vector H is mapped by the decoder to a vector X’ of dimension N that approximates the input vector X.

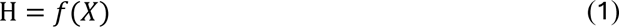

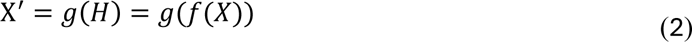

Vanilla autoencoders consist of an input layer, an output layer of size equal to the input size, and a hidden layer whose output is the latent representation of the data. When the hidden layer’s size equals or exceeds the input size, the autoencoder is called overcomplete. On the other hand, when the hidden layer’s size is smaller than the input size, the autoencoder is called undercomplete. Furthermore, when an autoencoder has more than one hidden layer, i.e., the encoder and decoder are deep neural networks, the autoencoder is called deep autoencoder.

Several variations of autoencoders have been proposed to prevent these from learning the identity function (especially in overcomplete autoencoders). Sparse autoencoders, for example, are vanilla autoencoders that use L1 regularization or the KL-divergence as a regularization term in the loss function to promote the sparsity of the model. Denoising autoencoders (Vincent et al., 2008) add noise to the input data and are trained to reconstruct and remove the noise of the data. Contractive autoencoders (Rifai et al., 2011) are trained to be robust to small variations in the input data by adding a penalty to the loss function. Usually, the penalty is the Frobenius Norm of the Jacobian matrix of the encoder activations for the input data.

### 2.3 Methodology

To classify the beehive sound signals, the audio files are divided into frames and a cluster analysis is performed over encoded representations of their acoustic features. The process preceding the cluster analysis is divided into two phases. In the first phase (called “*feature extraction process”*), the audios are divided into frames from which their MFCC-based features are extracted following the procedure proposed in (Scarpiniti et al., 2021b). In the second phase (called “*feature encoding process”*), the resulting dataset is split into a training and a test set at an 80:20 ratio. Then, the training set is used to train an artificial neural network model, first in an unsupervised manner as an autoencoder and then in a supervised manner as a classifier to learn a new representation of the audio frames. The feature extraction and feature encoding processes are discussed in more detail in the section 2.4. Each data point is labeled according to the audio file from which it is extracted; thus, 3 classes are considered in this paper, namely “100m”, “300m” and “500m”. The feature encoding process was conducted multiple times using different models with different depths and numbers of parameters to analyze how such properties affect the quality of the latent representation. Once the feature encoding process is completed, the test set is used to evaluate the features generated by the models using three metrics: Silhouette score, F1-score, and Accuracy.

The Silhouette score is a widely used metric that measures the quality of the clusters in terms of consistency; hence, it was used to evaluate the quality of the features generated. The Silhouette score for a data point (in this case a data point is an audio frame) is given by Equation (*3*):

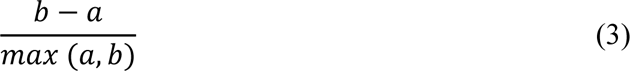

Where *a* is the mean distance between a data point and all other data points that belong to the same cluster (intra-cluster distance) and *b* is the mean distance between a data point and all other data points that belong to the neighboring cluster (the nearest cluster from which the data point is not a member). To calculate the Silhouette score of the whole test set, the mean of the Silhouette score over each one of the data points in the test set is used.

Accuracy and F1-score are metrics widely used to evaluate a classifier’s performance. Accuracy is a straightforward metric defined as the ratio of the correctly classified data points to the total number of points classified. The F1-score (Equation (4)), on the other hand, is defined as the harmonic mean of Precision and Recall. These two metrics are defined for binary classification tasks and are calculated following Equations (5) and (6), where TP is the number of true positives i.e., the number of data points correctly classified as belonging to the positive class, FP is the number of false positives i.e., the number of data points misclassified as belonging to the positive class, and FN is the number of false negatives i.e., the number of data points misclassified as belonging to the negative class. For multiclass classification tasks (such as the one tackled in the present paper), these metrics are calculated for each one of the classes, treating each one of them as the positive class and the rest as the negative class. Then, the results obtained are aggregated by adding them or averaging them. These metrics reach their best value at 1 and worst value at 0.

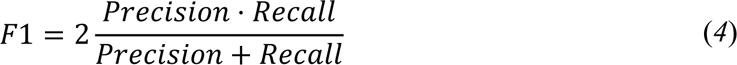

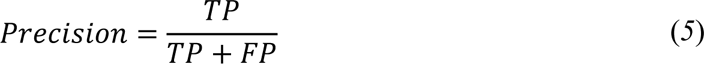

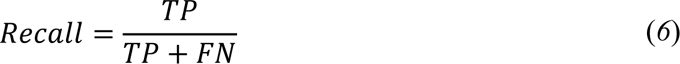

To further assess the quality of the features generated following the approach proposed in (Scarpiniti et al., 2021b), we also applied the K-means algorithm (MacQueen, 1967) in the raw feature space of the dataset (i.e., using the features obtained through the feature extraction process without the encoding process) and computed the Silhouette score values for the resulting clusters. To visually analyze the features generated, the Uniform Manifold Approximation and Projection (UMAP) algorithm (McInnes et al., 2018) was used to reduce their dimensionality, transforming them from their *H*_*s*_-dimensional space to a two-dimensional space. The UMAP is a fast and efficient general-purpose algorithm used for dimensional reduction based on manifold learning that has already been used to visualize and analyze features in machine learning-related works (Di et al., 2023; Mohammadimanesh et al., 2019; Zhang et al., 2021).

### 2.4 Data features

As mentioned in the previous section, the process preceding the cluster analysis is divided into two phases, the *Feature extraction process* and the *Feature encoding process.* In the first phase, the audio clips are divided into frames from which six statistics of their MFCCs are extracted. In the second phase, an Artificial Neural Network (ANN) is used to encode the MFCCs’ statistical data into a more meaningful representation of lower dimensionality. Although MFCC features-based approaches have already been used in beekeeping-related works (Di et al., 2023; Ribeiro et al., 2021; Soares et al., 2022; Terenzi et al., 2019; Zgank, 2018, 2022), the use of autoencoders for the analysis of sound signals in this research field remains mostly unexplored, with few exceptions (Cejrowski & Szymański, 2022; Davidson et al., 2020). To the best of our knowledge, MFCC feature-based approaches and autoencoders have not been used together in beekeeping-related works. It is important to emphasize that the main goal of the current paper is not to test the performance of the proposed approach against others on audio classification, but rather to determine through a cluster analysis whether the beehive sound contains spatial cues of the colony’s foraging sites.

#### 2.4.1 Feature extraction process

Following the feature extraction process proposed in (Scarpiniti et al., 2021b), each audio clip is first divided into *N*_*F*_ frames, following the Equation (7), where *C*_*D*_ is the clip duration and *S*_*F*_ is the frame size.

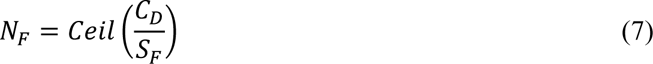

Since the sampling frequency for our audio clips is the same as that of the audio recordings analyzed in (Scarpiniti et al., 2021b) (*F*_*S*_ = 44,100*HZ*), all parameters for the feature extraction process were set the same. Hence, the frame size *S*_*F*_ = 100*ms*, which for our audio clips resulted in the number of frames reported in Table 2. In case the audio clip cannot be divided into equal size frames, the sample padding technique proposed in (Nguyen & Pernkopf, 2020) is used, filling the missing samples in the frames that need to be expanded with random samples taken from the same frame.

**Table 2.** Number of frames obtained for each class. After the feature extraction process, each frame is converted into a single data point with 384 features.

| Class | Frames - Data points |
| --- | --- |
| 100m | 24,219 |
| 300m | 25,616 |
| 500m | 24,853 |
| **Total** | **74,688** |

Each audio frame is further divided into smaller chunks from which a vector of 64 MFCCs is obtained. Considering a window size and a hop size of *S*_*W*_ = 2,048 and *S*_*H*_ = 512 respectively, and that each frame contains 4,410 samples, 9 vectors of 64 MFCCs are generated per frame. The first four statistical moments and the maximum and minimum of each MFCC are computed. As a result, each frame corresponds to a data point formed by a set of 64 × 6 = 384 statistical features. Figure 3 shows a graphical representation of the feature extraction procedure.

**Figure 3.**
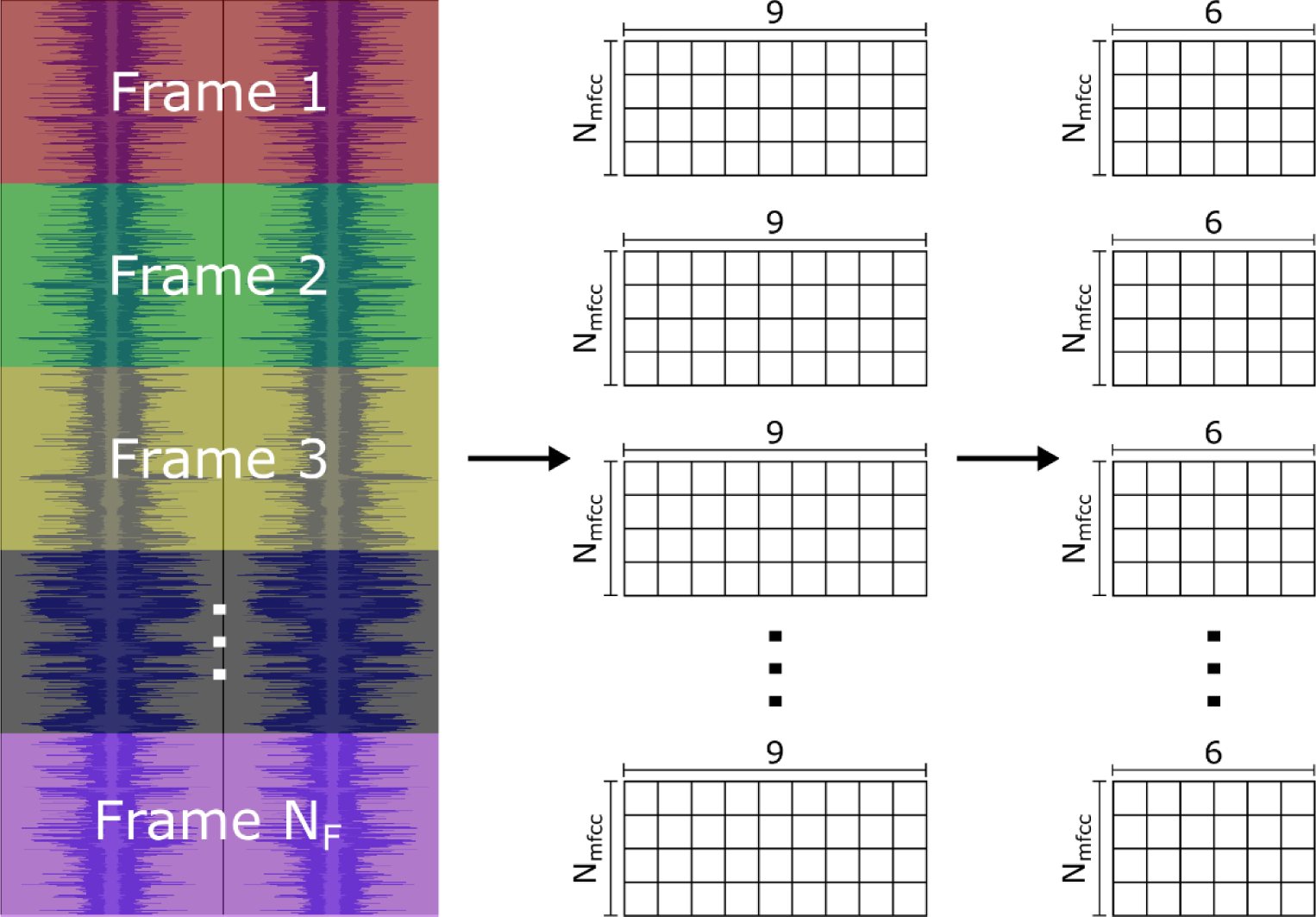
Graphical representation of the procedure followed to extract the features from the audio files. From left to right, first, the file is divided into N_F_ frames, then the 9 vectors of MFCCs are calculated for each frame. Finally, the maximum value, minimum value, and the first four statistical momentums are calculated for each MFCC, resulting in the 384 statistical features that represent each frame.

#### 2.4.2 Feature encoding process

In the feature encoding process, an ANN encodes the MFCCs’ statistical data into a new representation of lower dimensionality. The dataset obtained during the feature extraction process is divided into a training and a test set at a ratio of 80:20. As can be seen in Table 2, the three classes have a similar number of data points, which results in a balanced dataset that provides more confidence about the metrics used to evaluate the models.

The encoding process is divided into two stages, the pre-training stage, and the fine-tuning stage. In the pre-training stage, the training set is used to train a deep autoencoder in an unsupervised manner to reconstruct the MFCCs’ statistical data. After that, the decoder is detached from the pre-trained autoencoder, and a classification layer is added at the end of the encoder. Then, the resulting classifier is fine-tuned in a supervised manner on the same training set used in the pre-training stage.

It has been shown that pretraining classifiers as autoencoders allows them to learn a compressed and meaningful representation of data (Lopez Pinaya et al., 2020; Tschannen et al., 2018). This is a very convenient property for our classification problem, given that the MFCC-based features alone were not informative enough to observe any relevant differences between frames from different classes. Furthermore, as has been shown in (Erhan et al., 2010), unsupervised pre-training not only helps to initialize the model parameters in more promising areas within the parameter space but also helps to capture dependencies between parameters, resulting in better generalization which is also robust to random initialization. For this work, we considered three classes, one for each audio clip and distance to the hive. Hence each data point was labelled either 100m, 300m, or 500m. Additionally, the feature extraction procedure was conducted multiple times using different autoencoders with different depths and numbers of parameters to analyze how such properties affect the quality of the latent representation obtained.

## 3 RESULTS

A group of 16 models were evaluated to find a suitable topology for the autoencoder-classifier network. The configurations tested are listed in Table 3, where the column to the left shows the ID number of the model, the column at the center the configuration of each model during the pretraining phase (i.e. as an autoencoder), and the column to the right the final shape of the model after the pre-training, where the decoder is detached and a classifier layer is attached to it. Each element in the lists corresponds to a network layer, with the numbers being the number of neurons in that layer; “dropout” and “norm” indicate either a dropout layer or a batch normalization layer respectively, and *H*_*s*_ is the embedding size, which for each model ID took the values of 8, 16, 32 and 64. Models 1 and 2 differ solely on the dropout layer, and so do Models 3 and 4.

**Table 3.** Structure of the models tested to encode the MFCC-based features. The “Model ID” column shows the number ID of the models, and the “Shape” columns show the layer configuration of the models, both as autoencoders and classifiers. Here “dropout” and “norm” indicate either a dropout or a batch normalization layer. *H*_*s*_ is the embedding size with four possible values: 8, 16, 32, and 64. Hence, each model ID represents 4 different configurations.

| Model ID | Pre-training shape | Final shape |
| --- | --- | --- |
| 1 | [384, 384, dropout, 192, 192, norm, $H_s$, 192, 192, dropout, 384, 384] | [384, 384, dropout, 192, 192, norm, $H_s$, 3] |
| 2 | [384, 384, 192, 192, norm, $H_s$, 192, 192, 384, 384] | [384, 384, 192, 192, norm, $H_s$, 3] |
| 3 | [384, dropout, 192, norm, $H_s$, 192, dropout, 384] | [384, dropout, 192, norm, $H_s$, 3] |
| 4 | [384, 192, norm, $H_s$, 192, 384] | [384, 192, norm, $H_s$, 3] |

Three metrics were computed to evaluate the performance of the candidate models, namely Silhouette, F1-score, and Accuracy (see section 2.3). Since each model in its final form can be divided into encoder and classifier, the Silhouette score was used to evaluate the encoders, while the F1-score and Accuracy metrics were used to evaluate the classifiers. Additionally, the F1-score and Accuracy metrics further validate the quality of the encoded features, since high-quality features ease the classification.

### 3.1 Performance of autoencoder-classifier models

The performance metrics for the experiments run on the 16 proposed models are shown in Table 4 to Table 7. First, the models were trained using the training set. Then they were fed with the test set to compute the Silhouette score, F1-score, and Accuracy. Each experiment consisted of 30 instances per model to reduce the probability of spurious results. The values shown are the mean over the results obtained from the 30 instances of each model.

**Table 4.** Silhouette score, F1-score, and Accuracy of the outcomes of Model 1 when the embedding size was set to 8, 16, 32, and 64.

| Model 1 |  |  |  |  |
| --- | --- | --- | --- | --- |
| $H_s$ | 8 | 16 | 32 | 64 |
| **Silhouette score** | 0.927252 | 0.89967 | 0.840720 | 0.767208 |
| **F1-score** | 0.989135 | 0.98932 | 0.990225 | 0.989970 |
| **Accuracy** | 0.989135 | 0.98932 | 0.990226 | 0.989972 |

**Table 5.** Silhouette score, F1-score, and Accuracy of the outcomes of Model 2 when the embedding size was set to 8, 16, 32, and 64.

| Model 2 |  |  |  |  |
| --- | --- | --- | --- | --- |
| $H_s$ | 8 | 16 | 32 | 64 |
| **Silhouette score** | 0.819736 | 0.80023 | 0.747045 | 0.677263 |
| **F1-score** | 0.987631 | 0.990966 | 0.991257 | 0.991196 |
| **Accuracy** | 0.988225 | 0.990967 | 0.991259 | 0.991197 |

**Table 6.** Silhouette score, F1-score, and Accuracy of the outcomes of Model 3 when the embedding size was set to 8, 16, 32, and 64.

| Model 3 |  |  |  |  |
| --- | --- | --- | --- | --- |
| $H_s$ | 8 | 16 | 32 | 64 |
| **Silhouette score** | 0.782154 | 0.742183 | 0.686134 | 0.622511 |
| **F1-score** | 0.983291 | 0.984986 | 0.985387 | 0.985008 |
| **Accuracy** | 0.983286 | 0.984989 | 0.985393 | 0.985016 |

**Table 7.** Silhouette score, F1-score, and Accuracy of the outcomes of Model 4 when the embedding size was set to 8, 16, 32, and 64.

| Model 4 |  |  |  |  |
| --- | --- | --- | --- | --- |
| $H_s$ | 8 | 16 | 32 | 64 |
| Silhouette score | 0.690155 | 0.667827 | 0.608412 | 0.551401 |
| F1-score | 0.986511 | 0.988620 | 0.988950 | 0.990300 |
| Accuracy | 0.986638 | 0.988644 | 0.989182 | 0.990300 |

Additionally, to validate the effect of pre-training the models as autoencoders, a network with the same “final shape” as Model 1 (the model with the highest Silhouette score) was also included in the study. Following the notation used in Table 3, this network had shape [384, 384, dropout, 192, 192, norm, *H*_*s*_, 3] and will be referred to as “Model 5”. This model did not go through the pre-training stage as an autoencoder, and its performance metrics are shown in Table 8.

**Table 8.** Silhouette score, F1-score, and Accuracy of the outcomes of Model 5 when the embedding size was set to 8, 16, 32, and 64.

| Model 5 |  |  |  |  |
| --- | --- | --- | --- | --- |
| $H_s$ | 8 | 16 | 32 | 64 |
| Silhouette score | 0.916805 | 0.879652 | 0.823654 | 0.777524 |
| F1-score | 0.988831 | 0.989006 | 0.989400 | 0.989862 |
| Accuracy | 0.988829 | 0.989008 | 0.989403 | 0.989865 |

Figure 4 to Figure 8 show the feature space generated by the evaluated models. The features plotted were taken from an instance of the model type with the highest Silhouette score value (e.g., Figure 4 shows the features generated by an instance of Model 1 with *H*_*s*_ = 8). Each figure displays the ground truth classification of the data and the classification learned by the model along with the corresponding Silhouette score (notice that the Silhouette scores differ from those reported in the tables since the latter are average values over 30 instances).

**Figure 4.**
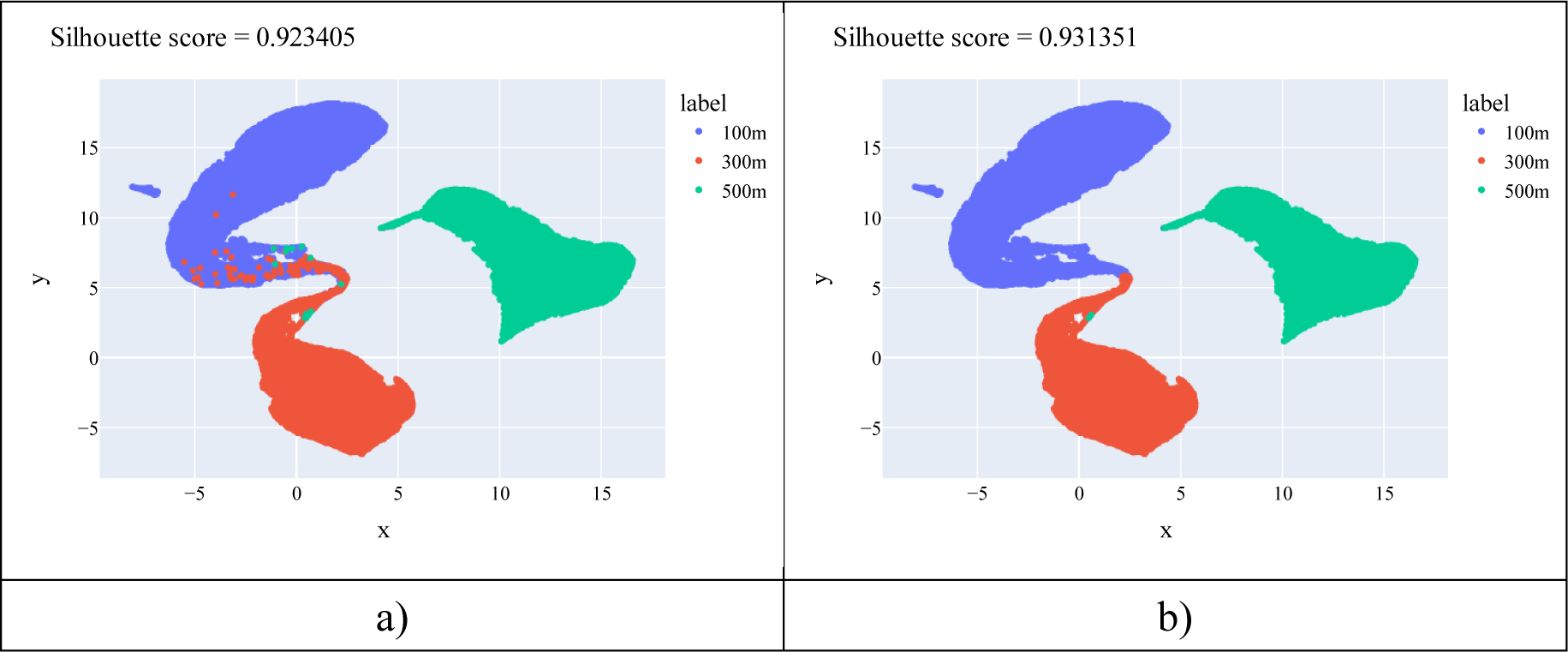
Feature space generated by Model 1. a) Shows the ground truth classification. b) Shows the classification given by the instance of the Model 1 which performance was closer to the average.

**Figure 5.**
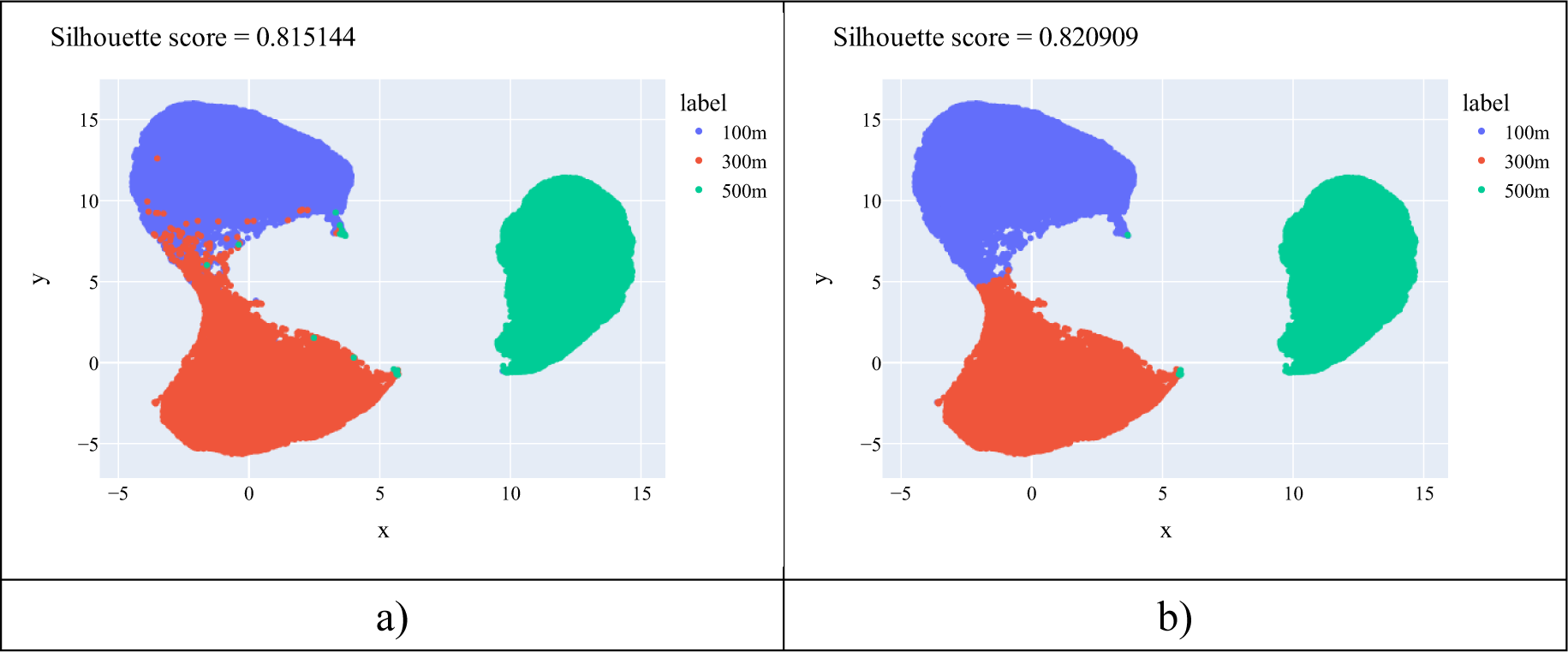
Feature space generated by Model 2. a) Shows the ground truth classification. b) Shows the classification given by the instance of the Model 2 which performance was closer to the average.

**Figure 6.**
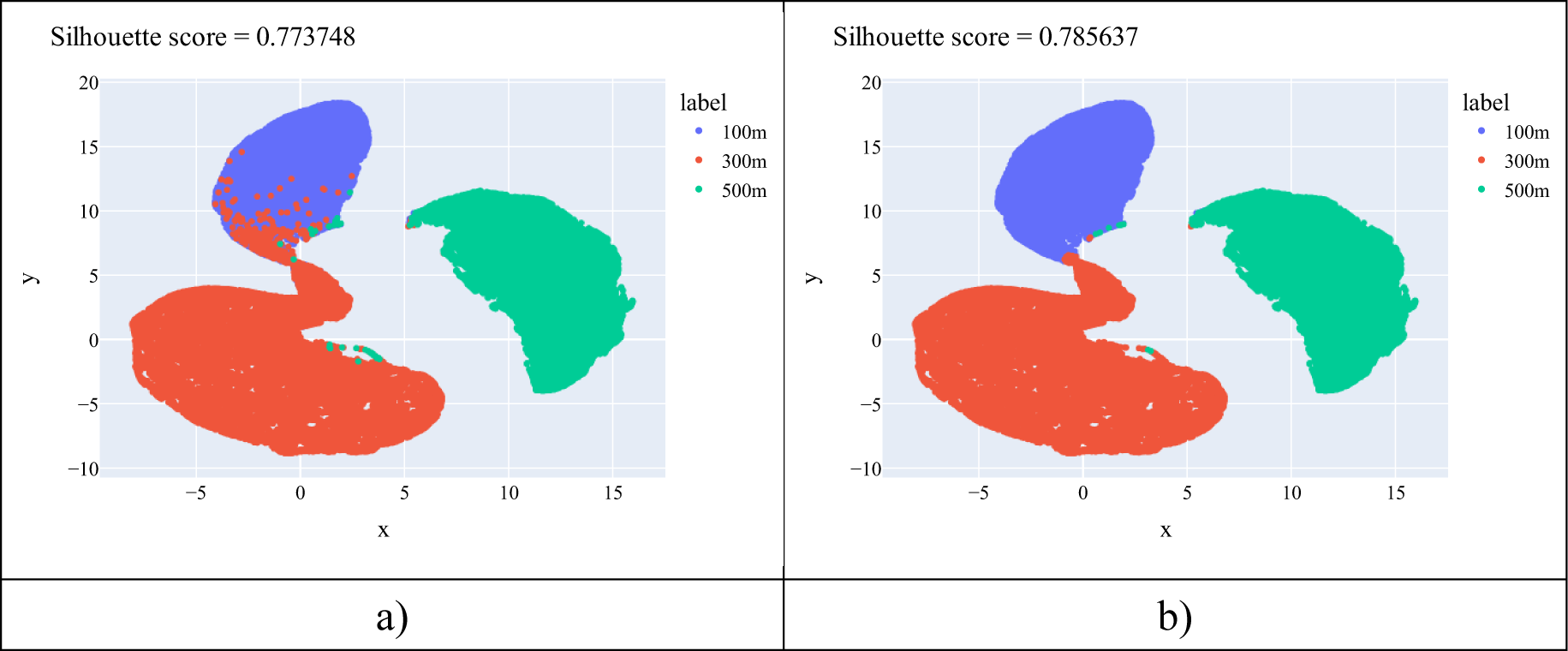
Feature space generated by Model 3. a) Shows the ground truth classification. b) Shows the classification given by the instance of the Model 3 which performance was closer to the average.

**Figure 7.**
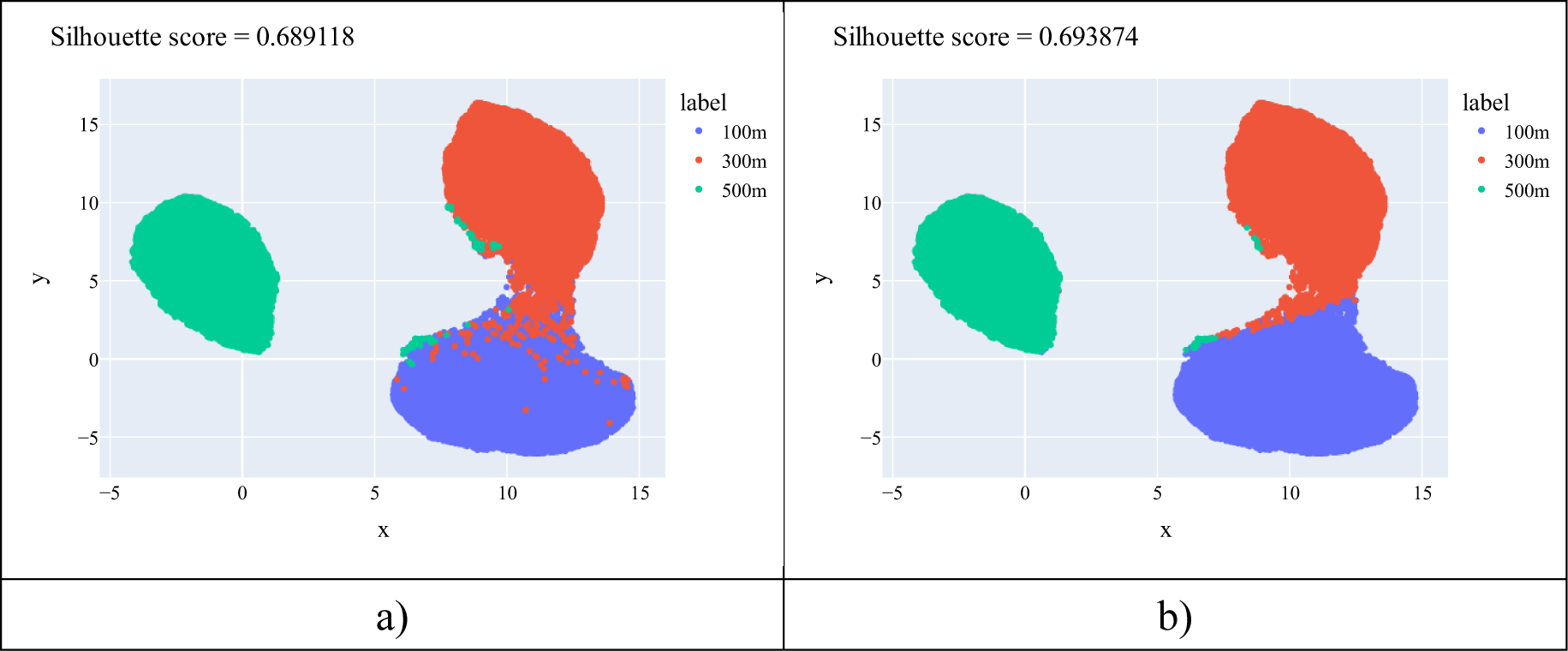
Feature space generated by Model 4. a) Shows the ground truth classification. b) Shows the classification given by the instance of the Model 4 which performance was closer to the average.

**Figure 8.**
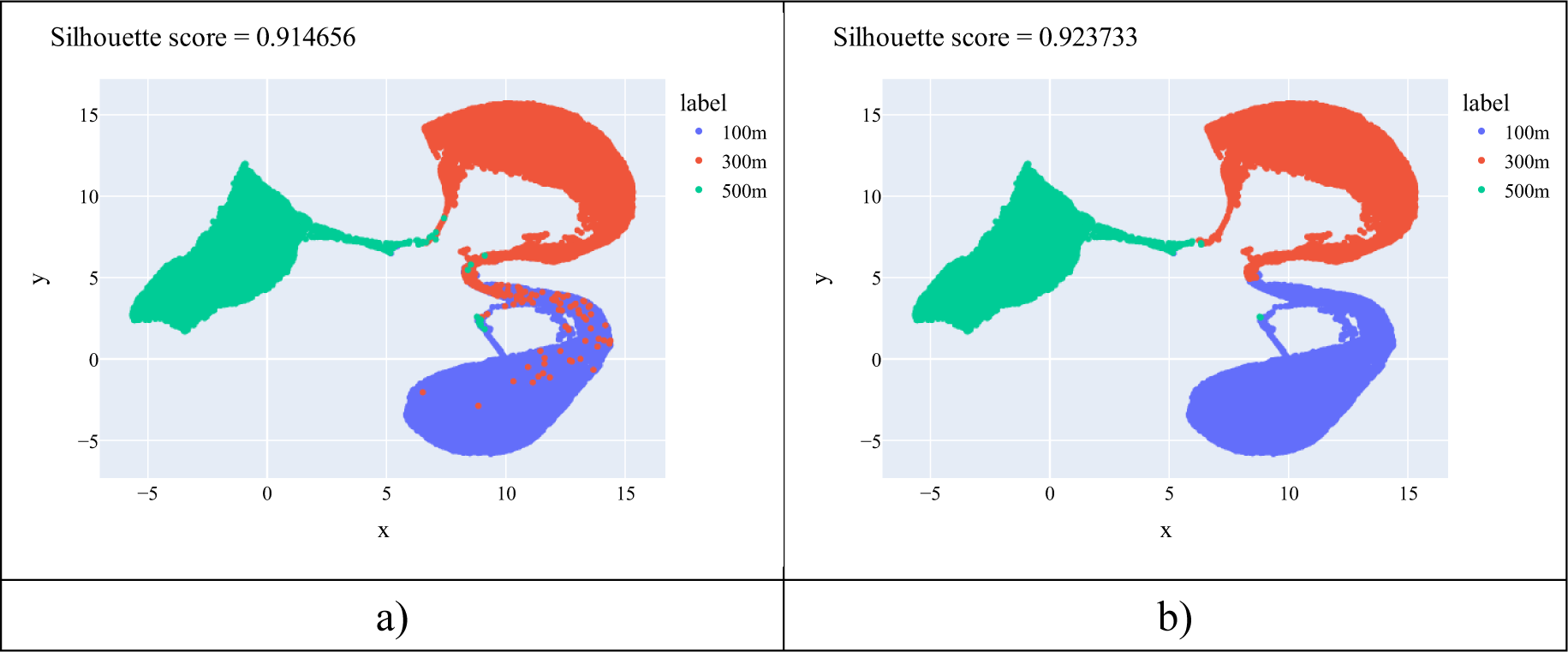
Feature space generated by Model 5. a) Shows the ground truth classification. b) Shows the classification given by the instance of the Model 5 which performance was closer to the average.

The classifications achieved by the models do not differ greatly from the ground truth classification, which is also reflected in the high Accuracy and F1-score. Models 1 and 3 obtained higher Silhouette scores than their versions without dropout (Models 2 and 4), evidencing that the dropout technique played a crucial role in learning richer features. Similarly, deeper models obtained better representations of the audio frames, as denoted by the higher Silhouette scores obtained by Models 1 and 2 compared to their shallow versions, Models 3 and 4 respectively. Furthermore, the feature spaces generated by Models 1 and 5 obtained the highest Silhouette scores in both class distributions, indicating the importance of both, the network structure and the pre-training stage, to obtain descriptive features that ease the audio classification.

Visually, Model 1 assigned features to the data points corresponding to audio 500m such that the corresponding cluster is completely isolated from the other two, while clusters formed by the data points of audios 100m and 300m show an area of intersection. By contrast, in the feature space generated by Model 5, the data points corresponding to audio 500m do not form an isolated cluster, instead, it has an area of intersection with the cluster formed by the data points corresponding to audio 300m. Besides, just like with Model 1 clusters formed by points from audios 100m and 300m have an intersection area, indicating that both are somehow related.

### 3.2 Performance of k-means clustering

To further validate the encoding process, we performed two experiments using the K-means algorithm to define the three clusters. In the first experiment, we applied the K-means algorithm to the raw feature space. Figure 9 shows the cluster assignations obtained by the K-means algorithm, the ground truth clusters, and their corresponding Silhouette score values on the raw feature space. From a visual analysis, it is evident that the clusters are not well-defined in the raw feature space, with data points of the three classes grouped in what could be interpreted as a single cluster, and only some of the 500m class data points isolated in a second cluster. This visual analysis is supported by the meager value of the Silhouette scores, proving that the raw features do not describe the data well enough to distinguish between the different classes.

**Figure 9.**
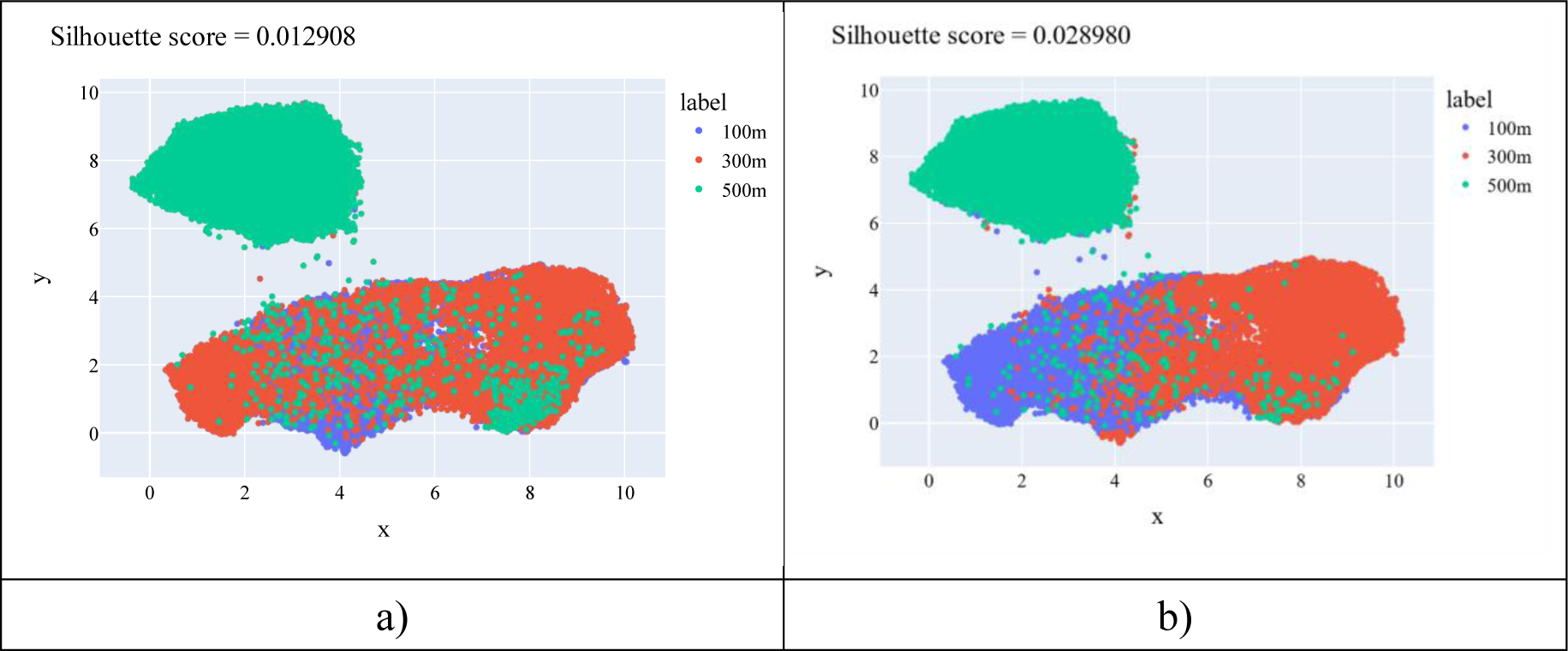
(a) Shows the ground truth labels on the raw feature space. (b) Shows the clusters identified by the K-means algorithm on the raw feature space.

In the second experiment, we used the latent space generated by the instance of Model 1 with *H*_*s*_ = 8 (the configuration with the highest Silhouette score). Figure 10 shows the cluster assignations obtained by the K-means algorithm, the ground truth clusters, and their corresponding Silhouette score values on this feature space. It is worth mentioning that, unlike the experiments with the autoencoder-classifier models, where the reported results only considered the test set, here we used the totality of the available data points (training and test set), hence the UMAP plot differs from the presented in Figure 4.

**Figure 10.**
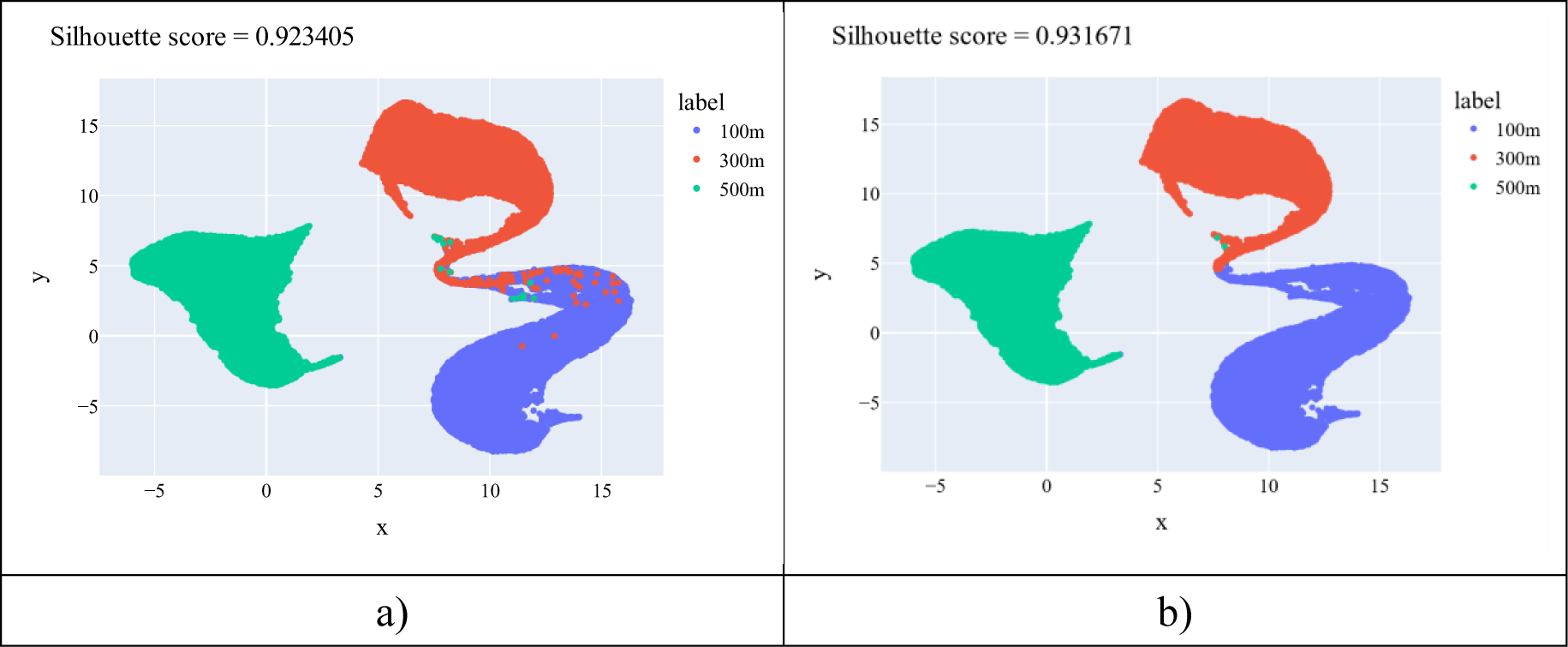
(a) and (b) show the ground truth labels on the feature space generated by Model 1 with Hs=8 and the clusters identified by the K-means algorithm, respectively.

From a visual analysis, it is clear that clusters in Figure 10 are better defined than those shown in Figure 9, which also results in significantly higher Silhouette scores, proving that the feature space generated by the autoencoder enhances the differences between classes. In further analysis of the feature space generated by Model 1, we can observe a well-defined cluster containing exclusively 500m data points. The clusters of classes 100m and 300m show a small overlapping, which can be interpreted as a certain resemblance between audios from these two classes. It is important to remember that the UMAP plots are projections of high-dimensional spaces into a 2-dimensional space. Hence, what appears as evident overlapping and intersection in the plots, might not be such in the original space.

## 4 DISCUSSION

Our results indicate sizable differences in the beehive sound of a colony when foraging from food sources located at different distances from the hive. Some of these differences are already visible in the feature space of the MFCC’s statistical features (see Figure 9), where most data points from the 500m class are isolated from those of classes 300m and 100m. However, the use of autoencoding networks greatly improved the quality of the feature space, resulting in better-defined clusters, including the separation of points corresponding to the classes 100m and 300m, previously grouped in a unique cluster in the raw feature space.

Interestingly, all evaluated models obtained the highest Silhouette score when the embedding size *H*_*s*_ was set to 8 (see Table 4 to Table 7) and not with a higher value. Also, for all models, the Silhouette score decreased as *H*_*s*_ increased, reaching the lowest value when *H*_*s*_ was set to 64. Nevertheless, the Silhouette score does not greatly affect the classifiers’ performance. As can be seen in Table 9, the average values of all the F1-scores and Accuracy were barely under 0.99 with small standard deviations, indicating a homogeneous performance of all models as classifiers. It is also worth noting that the highest values Accuracy and F1-score values were not reached with the lowest *H*_*s*_. The disagreement between the Silhouette scores and the clustering metrics suggests that the Silhouette score might not be the best choice when working with high-dimensional spaces.

**Table 9.** Mean and STD of the F1-scores and Accuracy obtained by Models 1, 2, 3, and 4.

|  | **F1-score** | **Accuracy** |
| --- | --- | --- |
| **Mean** | 0.988297 | 0.988359 |
| **SD** | 0.004515 | 0.003918 |

We also explored the effect of pre-training the models as autoencoders and then fine-tuned them as classifiers, obtaining slightly better results. Also, as the metrics indicate, deeper models overperformed their shallower counterparts, as was the case for models with and without a dropout layer.

In summary, using deep neural networks to analyze audio recordings of beehive sound during foraging activity, we have found a correlation between audio descriptors and the distance to the foraging sources for the first time. Although additional studies need to be conducted in this direction, these findings could be of great significance to further understand the different means of communication in honey bee colonies. Furthermore, the use of MFCC’s statistical features and autoencoders has proven to be a powerful tool for analyzing sound signals that could find applications beyond the scope of this study.

## ACKNOWLEDGEMENTS

The authors would like to express our gratitude to the Sericulture and Apiculture Research Institute of Yunnan Academy of Agricultural Sciences for providing the space and the colony to carry out the experiments and to their staff members Zhang Xuewen, Hu Zongwen, Zhao Hongmu, Miao Chunhui, Huang Xinqiu, and Zhou Chuntao for their valuable help.

## AUTHOR CONTRIBUTIONS

Conceptualization: N.D., F.L., and F.W.; methodology: N.D. and F.L.; software: D.B.; resources, supervision: F.L.; project administration: F.L. and F.W.; data curation: N.D., D.B., J.G., and F.W.; writing: D.B., J.G., F.L., N.D., and F.W.; visualization: D.B.

